# Crop diversity increases disease suppressive capacity of soil microbiomes

**DOI:** 10.1101/030528

**Authors:** Ariane L. Peralta, Yanmei Sun, Marshall D. McDaniel, Jay T. Lennon

**Author notes:** A.L.P and Y.S. contributed equally to this work. Abbreviations 2,4-diacetylphloroglucinol (DAPG); plant growth promoting rhizobacteria (PGPR); plant pathogen suppression (PPS); pyrrolnitrin (PRN).

## Abstract

Microbiomes can aid in the protection of hosts from infection and disease, but the mechanisms underpinning these functions in complex environmental systems remain unresolved. Soils contain microbiomes that influence plant performance, including their susceptibility to disease. For example, some soil microorganisms produce antimicrobial compounds that suppress the growth of plant pathogens, which can provide benefits for sustainable agricultural management. Evidence shows that crop rotations increase soil fertility and tend to promote microbial diversity, and it has been hypothesized that crop rotations can enhance disease suppressive capacity, either through the influence of plant diversity impacting soil bacterial composition or through the increased abundance of disease suppressive microorganisms. In this study, we used a long-term field experiment to test the effects of crop diversity through time (i.e., rotations) on soil microbial diversity and disease suppressive capacity. We sampled soil from seven treatments along a crop diversity gradient (from monoculture to five crop species rotation) and a spring fallow (non-crop) treatment to examine crop diversity influence on soil microbiomes including bacteria that are capable of producing antifungal compounds. Crop diversity significantly influenced bacterial community composition, where the most diverse cropping systems with cover crops and fallow differed from bacterial communities in the 1-3 crop species diversity treatments. While soil bacterial diversity was about 4% lower in the most diverse crop rotation (corn-soy-wheat + 2 cover crops) compared to monoculture corn, crop diversity increased disease suppressive functional group *prnD* gene abundance in the more diverse rotation by about 9% compared to monocultures. Identifying patterns in microbial diversity and ecosystem function relationships can provide insight into microbiome management, which will require manipulating soil nutrients and resources mediated through plant diversity.

## Introduction

Microbiomes are collections of microorganisms that live in close association with plants and animals. Certain microorganisms can confer benefits because they contain genes that aid in nutrient acquisition (Chaparro et al. 2012, Berendsen et al. 2012) while other microorganisms can protect hosts by preventing colonization by pathogens (Latz et al. 2012, Schlatter et al. 2017). For example, soils harbor a diverse collection of microorganisms that affect the evolution and ecology of plant populations (Lau and Lennon 2012, van der Putten et al. 2013, 2016). Many soil microorganisms establish intimate associations with plant roots, which can result in enhanced plant growth through many mechanisms (Mendes et al. 2013, 2015). One important mechanism through which soil microorganisms increase plant performance and fitness is via disease suppression. In this case, a healthy and robust soil microbiome can serve as a first line of defense for plants against soil-borne pathogens within the resident soil microbial community (Mendes et al. 2013, van der Putten et al. 2016) either directly through antibiosis or parasitism, or indirectly through enhancing plant immune responses (Mendes et al. 2013).

Plant-soil feedback theory provides a framework for assessing the mechanisms and outcomes microbiome dynamics. More specifically, there are many ways soil microbiomes can be managed to influence soil pathogens. One way is through crop selection. Specifically, individual crops can affect pathogen populations by altering chemical, physical, or biological properties in their rhizosphere (Raaijmakers et al. 2009, Berendsen et al. 2012). Further, recent attention is being paid to ecological intensification of farms (Tilman et al. 2011), and one of the promising specific management practices under this strategy is to diversify farms by rotating crops (Smukler et al. 2010, Lin 2011). The colloquial use of the term “rotation effect” has a long history of agronomic research (Karlen et al. 1994), and its origins are from the overwhelming evidence that rotating crops increase crop yield (Liebman and Dyck 1993, Karlen et al. 1994). From a management or conservation perspective, crop rotations are not the traditional form of increasing biodiversity. At any given time, the species richness on a farm using crop rotations is often one (i.e., monoculture), but there is a diverse suite of biochemical inputs from crops planted at different times. There is mounting evidence that this form of ‘temporal biodiversity’ may provide some of the same beneficial ecosystem functions as traditional spatial biodiversity (Zak et al. 2003), such as carbon sequestration, pest control, and nutrient cycling (Ball et al. 2005, McDaniel et al. 2014b, Tiemann et al. 2015, Venter et al. 2016). Despite this frequently found “rotation effect”, the underlying mechanism(s) in support of crop rotations are largely unknown, but might be related to crop diversity promoting plant-pathogen-suppressing microorganisms.

Often, plant pathogen suppression (PPS) is associated with soil microbial communities that have the capacity to produce antimicrobial compounds. Specifically, antibiosis has been linked to disease suppressive capacity, whereby the abundance of antagonistic bacteria has been associated reductions in fungal pathogens through competitive inhibition (Weller et al. 2002, Haas and Défago 2005). For example, bacterial production of secondary metabolites 2,4-diacetylphloroglucinol (DAPG) and pyrrolnitrin (PRN) are two potent toxins known to suppress fungal pathogens in soils (Garbeva et al. 2004a, 2004b, Haas and Défago 2005). However, the extent to which abiotic and biotic factors influence the abundance of such microbes remains unclear. Abiotic factors (e.g., salt, moisture, nutrients) can limit the strength and alter the direction of plant-soil feedbacks (Bever et al. 1997, Mills and Bever 1998, Packer and Clay 2000, Kulmatiski et al. 2008). It has been argued that edaphic features may be important or even required for PPS and might influence species interactions. In addition, aboveground features such as plant diversity could influence PPS. Specifically, plant diversity could increase the total soil bacterial diversity giving way to the “sampling effect” where species-rich ecosystems contain species that function at high levels (e.g., Tilman et al. 2002, Naeem and Wright 2003). Specifically, plant diversity could increase the probability of harboring PPS in the soil microbial community. Alternatively, plant diversity could modify soil microbial communities without influencing total diversity but rather through selecting for microorganisms that perform certain functions such as disease suppression. Some evidence suggests that PPS microorganisms are influenced by competition for iron, antibiosis, lytic enzymes, and induction of systemic resistance with host plant (Doornbos et al. 2012). For example, antibiosis has been linked to disease suppressive capacity, whereby the abundance of antagonistic bacteria has been associated with reductions in fungal pathogens through competitive inhibition (Weller et al. 2002, Haas and Défago 2005). Therefore, the abundance of PPS microbes may be a reflection of the total diversity of the soil microbial community, but this hypothesis has not been rigorously evaluated.

Given the unknown effect of crop diversity on PPS, we used a long-term (12 y) crop rotation study at the Kellogg Biological Station LTER to examine the effect of crop diversity on soil bacterial biodiversity and PPS potential. Specifically, our study addresses the following questions: (1) what is the relationship between crop diversity and soil microbial community composition and PPS? and (2) what is the role of changes in soil physicochemical properties on the crop diversity effect on soil microbial community composition, and PPS? We hypothesized that increased crop diversity would increase the diversity of the soil microbial community, and also increase the PPS in the soil through supporting a higher proportion of disease suppressive microbial taxa.

## Methods

### Site description and experimental design

We collected soils from the Biodiversity Gradient Experiment (<u>http://lter.kbs.msu.edu/research/long-term-experiments/biodiversity-gradient/</u>) at W.K. Kellogg Biological Station Long-Term Ecological Research (KBS LTER) site in southwest, Michigan, USA. Mean annual temperature is about 10 °C and mean annual precipitation is about 1000 mm yr^-1^ (Robertson and Hamilton 2015). The soils are Kalamazoo (fine-loamy) and Oshtemo (coarse-loamy) mixed, mesic, Typic Hapluadalfs formed under glacial outwash (Crum and Collins 1995). The crop rotation treatments at the Biodiversity Gradient Experiment included: monoculture corn (*Zea mays*, mC), corn with 1 red clover (*Trifolium pretense* L.), cover crop (C_1cov_), corn-soy *(Glycine max,* CS), corn-soy-wheat (*Triticum aestivum,* CSW), CSW with red clover (CSW_1cov_), CSW with red clover and cereal rye (*Secale cereal* L., CSW_2c_o_v_), and a spring fallow treatment that was just plowed every spring but contains 7-10 naturally-occurring plant species in the region (Table 1). This spring fallow treatment is considered the benchmark for plant diversity in the region, and under same tillage. Plantings of cover crop were dependent on the main crop in rotation (Smith and Gross 2006, 2007). The experiment was in a randomized complete block design, which included four blocks or replicates of each treatment. All plots received the same tillage at 15 cm depth, and no fertilizer or pesticides were applied to these plots.

**Table 1.** Cropping diversity treatments at the Kellogg Biological Station Long-term Ecological Research (KBS LTER) Biodiversity Gradient Experiment Plots. Plant treatments were established in 2000. Treatments were composed of monoculture, two-crop rotation, three-crop rotation +/-cover crops, and fallow plots (early successional) and soil collected during the corn phase of the rotation. Treatment abbreviations are in parentheses.

| Crop diversity treatment description | Number of crop species |
| --- | --- |
| (1) Continuous monoculture (mC) | 1 |
| (2) Continuous monoculture, one cover crop ($C_{1cov}$) | 2 |
| (3) Two-crop rotation (CS) | 2 |
| (4) Three-crop rotation (CSW) | 3 |
| (5) Three-crop rotation, one cover crop ($CSW_{1cov}$) | 4 |
| (6) Three-crop rotation, two cover crops ($CSW_{2cov}$) | 5 |
| (7) Spring Fallow/early successional field (fallow) | 10 |

### Soil sampling

We sampled soil from six crop diversity treatments, but to eliminate any immediate crop effect all the treatments were sampled in the corn phase and a spring fallow treatment (Table 1) on November 1, 2012. In each plot, we collected five soil cores (5 cm diameter, 10 cm depth) and then homogenized the cores in the field. A subsample from each composite sample was sieved through 4 mm in the field, flash frozen in the field in liquid nitrogen, and stored at -80 °C prior to molecular-based microbial analyses.

### Soil physicochemical analyses

From the same soil samples that were flash frozen for DNA extraction, soil chemical properties (total carbon, total nitrogen, ammonium, nitrate, pH, texture). These soils were previously analyzed and soil physicochemical characteristics reported (McDaniel et al. 2014a, McDaniel and Grandy 2016). Labile C was measured as permanganate oxidizable C (POXC) according to (Culman et al. 2012). Overall biological activity and amount of potentially mineralizable carbon (PMC) and nitrogen (PMN) were analyzed using a 120 d aerobic incubation (McDaniel and Grandy 2016).

### Bacterial community sequencing

To examine the relationship between crop diversity and soil microbial diversity, we used 16S rRNA targeted amplicon sequencing of the soil bacterial community. We extracted DNA using the MoBio Power Soil DNA Isolation Kit (MO BIO Laboratories, Inc., Carlsbad, CA). DNA concentration was adjusted to a standard concentration of 20 ng μl^−1^ and used as template. To characterize bacterial taxonomic diversity, we used barcoded primers (515f/806r primer set) developed by the Earth Microbiome Project to target the V4-V5 region of the bacterial 16S subunit of the ribosomal RNA gene (16S rRNA) (Caporaso et al. 2012). For each sample, PCR product combined from three 50 μl reactions, concentration quantified, and PCR product from each soil sample was combined in equimolar concentrations for paired-end 250×250 sequencing using the Illumina MiSeq platform according to details in (Muscarella et al. 2014). Briefly, we assembled the paired-end 16S rRNA sequence reads using the Needleman algorithm (Needleman and Wunsch 1970). All sequences were subjected to systematic checks to reduce sequencing and PCR errors. High quality sequences (i.e., >200 bp in length, quality score of >25, exact match to barcode and primer, and contained no ambiguous characters) were retained. In addition, we identified and removed chimeric sequence using the UCHIME algorithm (Edgar et al. 2011). We aligned our sequence data set with the bacterial SILVA-based bacterial reference database (Yilmaz et al. 2014). During data analysis, operational taxonomic units (OTUs) were binned at 97% sequence identity and phylogenetic classifications of bacterial sequences performed. Sequences were processed using the software package *mothur* v.1.35.1 (Schloss et al. 2009, Kozich et al. 2013). A total of 12,539,359 sequence reads were generated, and we analyzed 47,261 OTUs for bacterial community analyses.

### Composition and abundance of disease suppression genes

To characterize the subset of the microbiome associated with disease suppressive potential, we targeted disease suppressive taxa as the subset of soil microorganisms possessing genes that are required for the production of antifungal compounds 2,4-diacetylphloroglucinol (DAPG) (von Felten et al. 2011) (see supplemental material) and pyrrolnitrin (PRN) (Garbeva et al. 2004b, Haas and Défago 2005).

We assessed the relative abundance of disease suppressive functional genes by targeting *prnD* using quantitative PCR (qPCR) (Garbeva et al. 2004b). The partial *prnD* gene abundance was quantified using a SYBR green assay with primers prnD-F (5’-TGCACTTCGCGTTCGAGAC-3’) and prnD-R (5’-GTTGCGCGTCGTAGAAGTTCT-3’) (Garbeva et al. 2004b). The 25 μL PCR reaction contained 1× GoTaq Colorless Master Mix (Promega, Madison, WI), 0.4 μM of each primer, and 5 μL of template DNA. Cycling conditions were as following: initial cycle 95 °C for 10 min, and 30 cycles of 95 °C for 15 s and 60 °C for 1 min. For the qPCR standard curve, *prnD* gene was amplified from soil genomic DNA. PCR fragments were cloned to pGEM-T Easy Vector System according to the manufacturer’s manual (Promega, Madison, WI). Plasmids were extracted using the QIAprep Spin Miniprep kit (Qiagen, Valencia, CA), and cloned fragments were verified by PCR and agarose gel electrophoresis. Dilutions of plasmid DNA containing *prnD* gene were used to generate standard curves in quantities ranging from 5.03×10^2^ to 5.03×10^7^ copies. We quantified the *prnD* gene in 25 μL reaction volumes containing about 20 ng DNA template, 13×TaqMan Environmental Master Mix 2.0 (Applied Biosystems, Valencia, CA), 13×SYBR green I, and 0.4 μM of each primer. Fragments were amplified with an initial denaturation step at 95 °C for 10 min, followed by 40 cycles of 95°C for 15 s, 60 °C for 1 min. For each sample, PCR reactions were run in triplicate. We obtained standard curves based on serial dilutions of mixed PCR product amplified from soil samples. Reactions were analyzed on a BIO-RAD CFX-96Real-Time System (Bio-Rad, Hercules, California, USA).

### Statistical analyses

We examined microbiome differences among crop diversity treatments by comparing total community diversity and composition as well as disease suppression markers. We tested for differences in total bacterial diversity (based on Shannon Diversity Index H’, bacterial species richness, and Pielou’s Evenness Index J’) and *prnD* gene abundance in response to crop diversity treatment using analysis of variance (ANOVA). We checked that data met assumptions of analyses, and we treated crop diversity treatment as a fixed factor and block as a random effect. We used Tukey’s Honestly Significant Difference (HSD) tests to identify between-group differences in bacterial diversity and *prnD* gene abundance.

To visualize patterns of microbial community composition, we used Principal Coordinates Analysis (PCoA) of the microbial community composition based on the Bray-Curtis dissimilarity coefficient for each possible pair of samples using the R statistical package (R Core Development Team 2015). To test for differences in total bacterial communities and a subset of previously identified biocontrol bacterial taxa (i.e., *Psuedomonas* spp. and *Streptomyces* spp.) among crop diversity treatments, we used non-parametric permutational multivariate analysis of variance (PERMANOVA) implemented with the *adonis* function in the R Statistics Package R version 3.2.3 (R Development Core Team 2015). PERMANOVA was also used to assess the contribution of soil factors to the variation in bacterial community composition. The R^2^ value reported refers to the treatment sums of squares divided by the total sums of squares for each soil factor in the model. Because the *adonis* function carries out sequential tests (similar to Type I sums of squares) (Oksanen et al. 2010), the effect of the last soil factor or soil biological activity factor of the model was included in the final PERMANOVA model summary (Peralta et al. 2012). We also performed a similarity percentage analysis (SIMPER) using the *simper* function (R Statistics Package R version 3.2.3) (Clarke 1993, Warton et al. 2012) to identify the bacterial OTUs responsible for community differences between monoculture corn and other crop diversity treatments and is based on the contribution of individual taxa to the average Bray-Curtis dissimilarity. We also performed multiple linear regression (gene abundance ~ crop number + total soil carbon + soil moisture + soil ammonium + soil nitrate) to test the influence of soil factors and crop diversity number on abundance of disease suppression/biocontrol gene *prnD* using the *lm* function in the R Statistics Package R version 3.0.2 (R Core Development Team 2015).

## Results

### Bacterial community composition and soil function relationships

The crop diversity treatment significantly influenced soil microbiomes represented by the bulk soil bacterial community composition (*R^2^* = 0.37, *p* < 0.001; Appendix S1: Table S2, Fig. 1). Bacterial communities from the fallow plots and the most diverse crop rotations (CSW, CSW_1cov_, CSW_2cov_) were more similar to each other than the lower crop diversity treatments (C_1cov_, CS) (Fig. 1). The monoculture corn (mC) treatment was more distinct in bacterial community composition than all other crop diversity treatments (Fig. 1).

**Figure 1.**
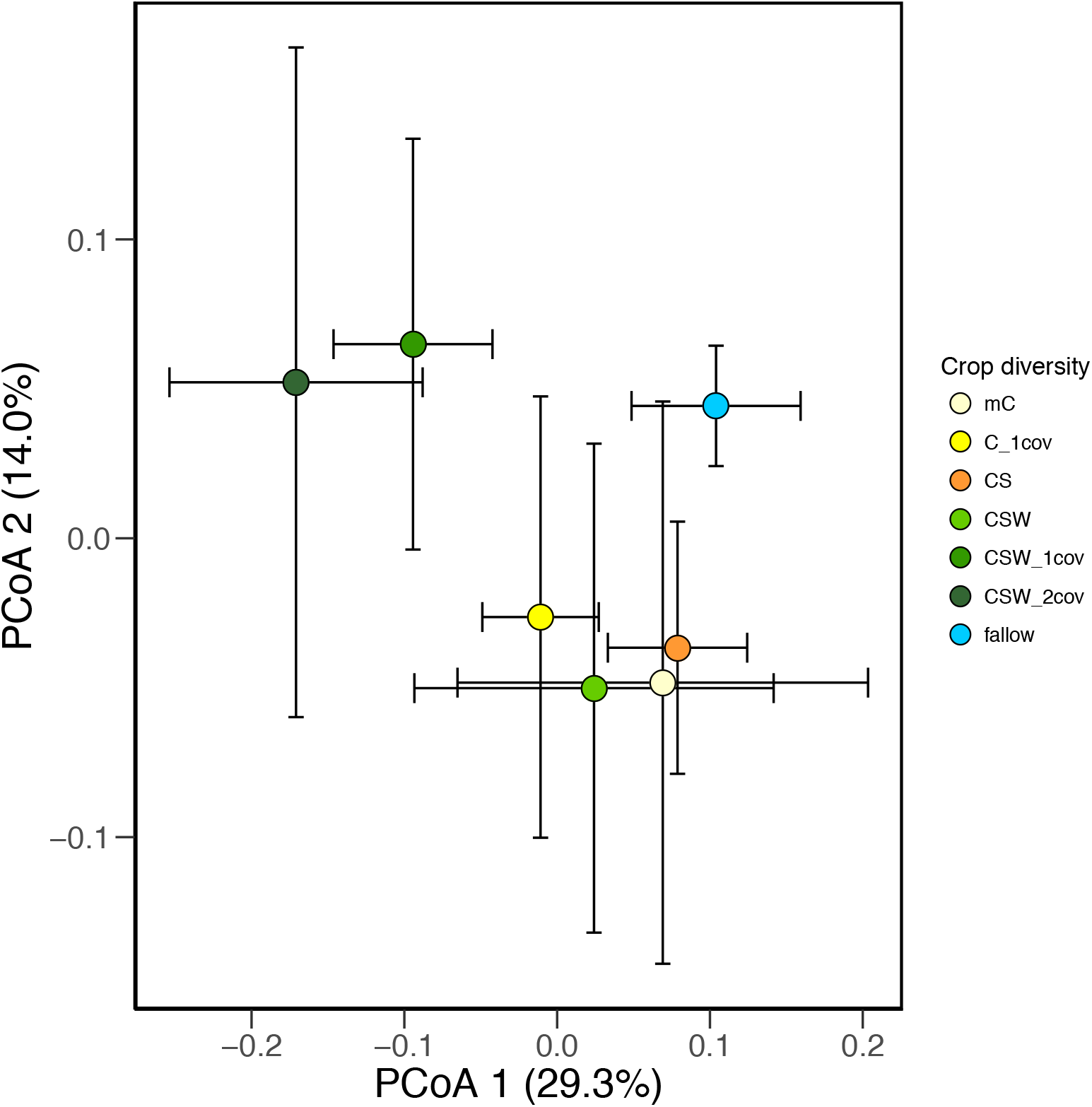
Ordination from Principal Coordinates Analysis depicting soil bacterial communities along a cropping diversity gradient. Symbols are colored according to cropping diversity treatment (mC=monoculture corn; C_1cov_ =corn/1 cover crop; CS=corn/soy; CSW=corn/soy/wheat; CSW_1cov_ =corn/soy/wheat/1 cover crop; CSW2_cov_=corn/soy/wheat/2 cover crops; fallow=spring fallow, tilled annually).

Bacterial diversity, as measured using Shannon Diversity Index (H’), was surprisingly greater under lower crop diversity systems than higher crop diversity systems, but highest in fallow treatments the most diverse non-cropping system (crop rotation: *F_6,20_*=10.16, *p*<0.0001; block: *F_1,20_*=0.20, *p*=0.6600; Fig. 2). Among, the corn cropping systems, mC had the highest Shannon Diversity Index in the most diverse rotation of corn-soybean-wheat with two cover crops (CSW_2cov_). In addition, bacterial species richness and Pielou’s Evenness Index (J’) revealed similar patterns across crop diversity treatments (evenness: *F_6,18_*=2.36, p=0.073; richness: *F_6,18_*=2.61, *p*=0.053; Fig. 2). Across all diversity metrics, the longest crop rotation (CSW_2cov_) showed the lowest richness and evenness values, and fallow soils generally had the highest values (Fig. 2).

**Figure 2.**
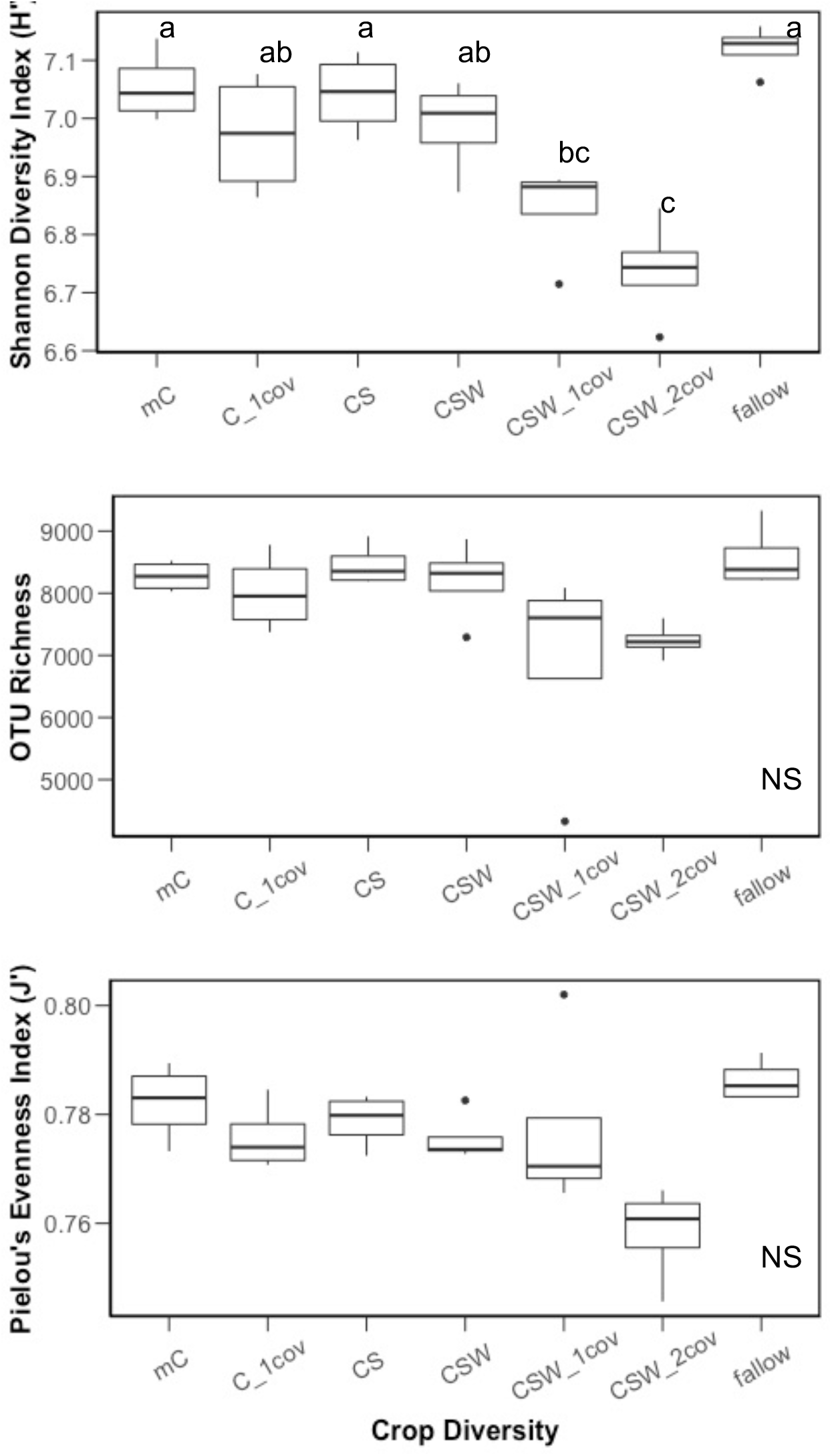
Total bacterial diversity (mean ± SEM based on Shannon Diversity Index H’), richness, and evenness (Pielou’s Evenness Index J’) in response to long-term crop diversity treatment. Different letters above points reflect significant differences in gene abundance along crop diversity gradient at *p* < 0.05 (Tukey’s HSD *post-hoc* analysis).

Soil physicochemical properties and soil function were related bacterial community composition to varying degrees. A summary of soil attributes is presented in Appendix S1: Table S1 and elsewhere (McDaniel and Grandy 2016). Bacterial community composition was best explained by soil texture, which varied across the experiment site from 9 to 38 *%* clay (R^2^=0.066, *p*<0.05, Table 3a). However, bacterial community composition was also marginally affected by soil moisture (R^2^=0.048, *p*<0.10, Table 2). Labile C as measured with permanganate oxidization was related to bacterial community composition (R^2^=0.074, *p*<0.05), but potentially mineralizable C did not. Potentially mineralizable nitrogen (PMN), however, which is produced in the same aerobic incubation as PMC and an indicator of nutrient-supplying power of a soil (a biologically available N pool), was significantly correlated with bacterial community composition (R^2^=0.063, *p*<0.05, Table 3).

**Table 2.** Summary of the contribution of (A) soil factors (original data from McDaniel et al. 2014) and (B) soil biological activity (original data from McDaniel and Grandy 2016) on bacterial community variation at the KBS Biodiversity Gradient Experimental Plots based on permutational MANOVA (PERMANOVA). Soil factor effects were considered to significantly contribute to community variation at *P* < 0.05.

| Effect | df | SS | MS | *F* | *R*<sup>2</sup> | *p*-value |
| --- | --- | --- | --- | --- | --- | --- |
| Sand | 1 | 0.088 | 0.088 | 2.243 | 0.066 | 0.014 |
| Silt | 1 | 0.088 | 0.088 | 2.239 | 0.066 | 0.020 |
| Clay | 1 | 0.087 | 0.087 | 2.207 | 0.065 | 0.024 |
| pH | 1 | 0.057 | 0.057 | 1.444 | 0.043 | 0.143 |
| Nitrate | 1 | 0.023 | 0.023 | 0.593 | 0.018 | 0.893 |
| Ammonium | 1 | 0.019 | 0.019 | 0.496 | 0.015 | 0.966 |
| Nitrogen | 1 | 0.043 | 0.043 | 1.086 | 0.032 | 0.326 |
| Carbon | 1 | 0.036 | 0.036 | 0.921 | 0.027 | 0.491 |
| Moisture | 1 | 0.064 | 0.064 | 1.622 | 0.048 | 0.078 |
| Residuals | 18 | 0.707 | 0.039 |  | 0.534 |  |
| Total | 27 | 1.325 |  |  | 1 |  |

(b) Soil Biological Activity
| Effect | df | SS | MS | *F* | *R*<sup>2</sup> | *p*-value |
| --- | --- | --- | --- | --- | --- | --- |
| PMN | 1 | 0.083 | 0.083 | 1.821 | 0.063 | 0.049 |
| PMC | 1 | 0.062 | 0.062 | 1.358 | 0.047 | 0.146 |
| POXC | 1 | 0.097 | 0.097 | 2.125 | 0.074 | 0.028 |
| Residuals | 24 | 1.100 | 0.046 |  | 0.830 |  |
| Total | 27 | 1.325 |  |  | 1 |  |

**Table 3.**
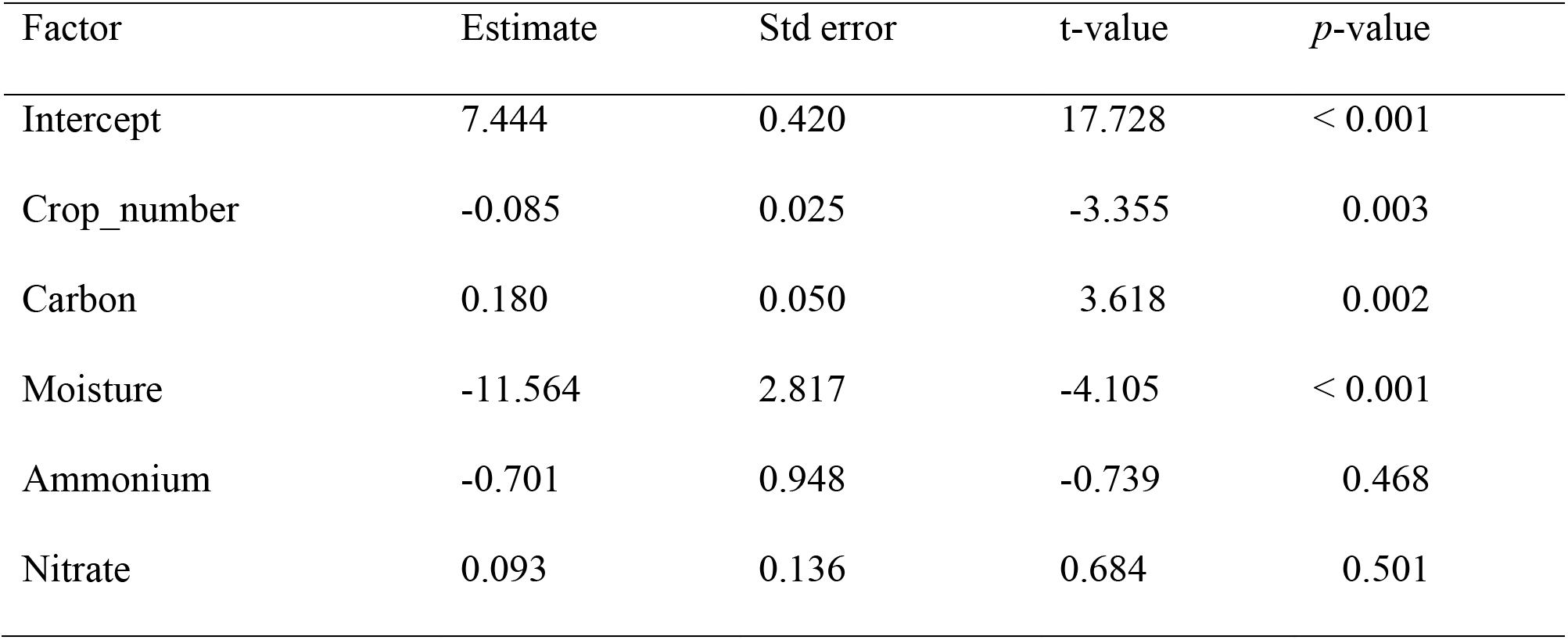
Summary of multiple linear regression to test the influence of disease suppressive functional potential (*prnD* gene abundance) on soil factors and crop diversity.

| Factor | Estimate | Std error | t-value | *p*-value |
| --- | --- | --- | --- | --- |
| Intercept | 7.444 | 0.420 | 17.728 | < 0.001 |
| Crop_number | -0.085 | 0.025 | -3.355 | 0.003 |
| Carbon | 0.180 | 0.050 | 3.618 | 0.002 |
| Moisture | -11.564 | 2.817 | -4.105 | < 0.001 |
| Ammonium | -0.701 | 0.948 | -0.739 | 0.468 |
| Nitrate | 0.093 | 0.136 | 0.684 | 0.501 |

### Disease suppression functional potential

Crop diversity affected PPS potential in soils. The *prnD* gene abundances in cropping systems were higher than under fallow conditions (crop rotation: *F_6,20_*=7.51, *p*=0.0003; Fig. 3). In cropping systems, the *prnD* gene in CSW_2cov_ treatment was the most abundant, and the gene abundance was significantly higher than in CSW and fallow treatments (Fig. 3). Our diversity benchmark, the fallow treatment (i.e., lowest crop diversity), showed the lowest *prnD* gene abundances (Fig. 3). Based on multiple linear regression analysis, plant and soil factors significantly related to *prnD* abundance (Adjusted R^2^=0.40, *F*=4.571, *p*=0.005). Crop species number (*p*=0.003), soil carbon (*p*=0.002), and soil moisture (*p*=0.0005) appeared to be significant predictors of *prnD* gene abundance (Table 3). We also observed a shift in the composition of disease-suppression microorganisms (represented by *phlD* gene fingerprint analysis using terminal restriction length polymorphism, T-RFLP) along the crop diversity gradient. The *phlD* community composition in the fallow treatment was different from other cropping systems (Appendix S1: Fig. S1).

**Figure 3.**
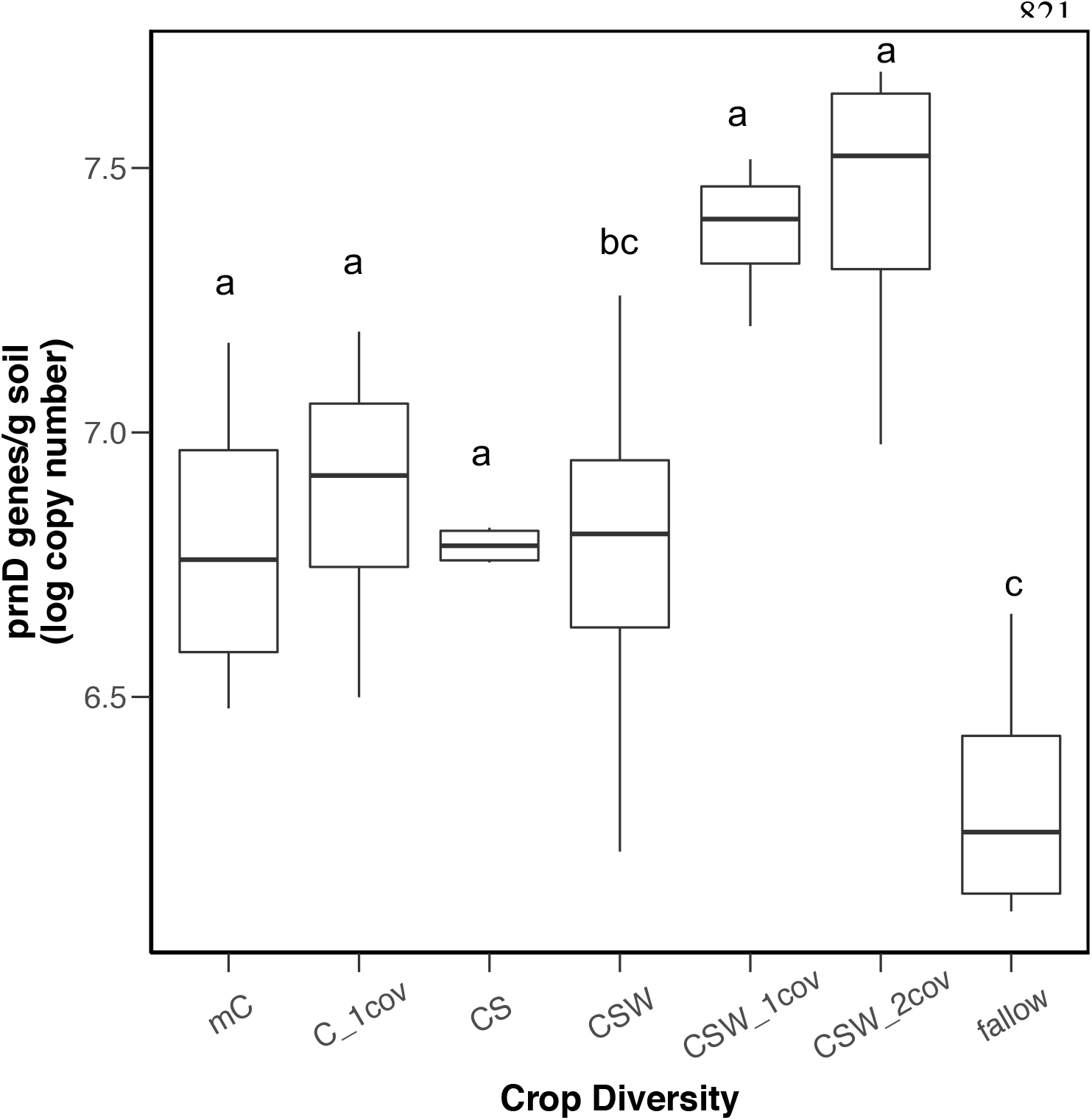
Abundance of prnD gene (PRN producers) in response to crop diversity treatment analyzed using quantitative PCR and expressed as log copy number of *prn*D gene. Different letters above points reflect significant differences in Different letters above boxplots considered significantly different in gene abundance at *p* < 0.05 (Tukey’s HSD *post-hoc* analysis).

### Soil bacterial-disease suppressive function relationship

The bacterial taxa primarily responsible for treatment differences between mC and the other crop diversity treatments were *Sphingomonadales* spp. and *Acidobacteria* subgroup Gp6 (Appendix S1: Table S3). When we compared a subset of taxa representing broad biocontrol bacterial community (composed of *Streptomyces* spp. and *Pseudomonas* spp.), there was no significant pattern in community composition across the crop diversity treatment (PERMANOVA; crop rotation: *R*^2^=0.321, *p*=0.132; Appendix S1: Table S4).

## Discussion

Soil microbiomes represent microbial communities living in close association with host plants and can protect host organisms from infection and disease. In this study, we found that crop rotation history impacted soil microbiomes and altered disease suppression potential in agricultural soils. However, we found some unexpected results that contrasted with our hypothesis. Contrary to our hypothesis, bacterial diversity decreased with increasing cropping diversity (Fig. 2). However, the PPS capability of the soil microbial community increased with crop diversity, but surprisingly the lowest PPS was in the diverse fallow treatments (Fig. 3). We observed that without crop plants (as reflected in the no crop fallow treatment), disease suppressive potential was significantly diminished compared to crop treatments, possibly due to reduced selection for soil microorganisms with disease suppression traits. The composition of the soil microbial community may be more important than diversity to soil suppressive function. Thus, crop rotation has the potential to impact diseases suppressive function, providing evidence for facilitation of fungal pathogen protection of plants in diverse crop rotation systems.

### Crop diversity effects on soil bacterial diversity

Crop rotation history decreased soil bacterial diversity over this 12-year crop diversity study. The pattern of reduced bacterial diversity (based on 16S rRNA gene sequencing) was lower in soils with higher cropping diversity. There are two most parsimonious explanations for this unexpected finding. First, this pattern in belowground biodiversity might be due to increased abundance of weedy plant species in low diversity treatments, but especially the monoculture corn. In other words, while we were considering the corn treatment as a single species; there could ostensibly have been up to 13 weed species per m^2^, as measured in an earlier study from this experiment (Smith and Gross 2007). On the other hand, this same study showed the most diverse cropping systems (CSW_2cov_) had only 5 or 6 weeds per m^2^. Second, perhaps there was not an artifact from the weeds and that soil bacterial diversity does decrease with increasing crop diversity, but other members of the soil microbial community (e.g., fungi, archaea) may be increasing in diversity with longer crop rotations. Despite decreased bacterial taxonomic diversity, a previous study based on the same soils that we used in this study, found that catabolic evenness (a measure of the diversity of catabolic function) also decreased with increasing crop diversity (McDaniel and Grandy 2016). This indicates that the trend in lower bacterial diversity with increasing crop diversity is not just structural, but also functional, and may indicate carbon resource specialization among bacteria since they are probably the major contributor to C catabolism in these substrate-induced respiration methods (Goldfarb et al. 2011, Allison et al. 2014). Based on phospholipid fatty acid analysis, a previous study showed that bacterial biomass in the micro-aggregate soil organic matter fraction was greatest in high compared low crop diversity treatments at this long-term experiment during a different sampling date (Tiemann et al. 2015). In addition, a previous meta-analysis revealed that the crop rotation effect increased soil bacterial diversity (i.e., Shannon Diversity Index H’) most notably in the first five years of treatment, but crop rotations longer than five years were more variable in diversity and not significantly different (Venter et al. 2016). Other studies do find significant negative effects of crop rotations on soil microbial diversity (Berg and Smalla 2009, Yin et al. 2010, Kulmatiski and Beard 2011, Reardon et al. 2014). The reason for these findings remain unknown but may be a combination of diversity impacts on other soil organisms not evaluated in this study or due to length of time associated with crop diversity treatment.

### Crop diversity and plant pathogen suppression relationship

We found that the increased crop diversity, via rotation, increased the abundance and altered the composition of a specific plant pathogen suppression gene (Figs. 3, Appendix S1: Fig. S1). Our results suggest that crop diversity may increase the disease suppression of agricultural soils, and are consistent with previous studies suggesting that plant diversity can enhance protection against soil-borne pathogens by fostering antagonistic soil bacterial communities (Latz et al. 2012, van der Putten et al. 2016). One potential explanation for the negative plant diversity and disease suppressive function relationship is due to facilitation, where changes in plant root exudation may lead to enrichment of plant growth promoting rhizobacteria (PGPRs) (Lugtenberg and Kamilova 2009, Badri et al. 2009, Chaparro et al. 2012). In previous studies, microbial interactions among the total microbial community and soil-borne pathogens in the plant rhizosphere have influenced both plant growth and productivity (Bakker et al. 2010, Penton et al. 2014).

The addition of cover crops to rotations strongly increased disease suppressive potential. This along with evidence from previous studies shows that crop rotations may prevent many forms of crop disease caused by *Fusarium* spp., *Phytophthora,* and *Rhizoctonia* spp. (Raaijmakers et al. 2009, van der Putten et al. 2016). Soil microbial diversity has been implicated as important for soil disease suppression; sterilized soils lose suppressive capacity, and adding soil microorganisms to sterilized soil facilitates disease suppression functional capacity (Garbeva et al. 2006, Brussaard et al. 2007, Postma et al. 2008). Biocontrol bacteria can also provide disease suppression against plant pathogens by way of the following mechanisms: competition for iron, antibiosis, lytic enzymes, and induction of system resistance of host plants (Doornbos et al. 2012, Schlatter et al. 2017). Plants can also facilitate recruitment of specific biocontrol microorganisms in some cases. A previous study suggests that beneficial pseudomonads are recruited depending on the most dominant soil-borne pathogen infecting crop species (Mavrodi et al. 2012, Berendsen et al. 2012). In the present study, we analyzed a subset of previously reported biocontrol bacterial taxa (e.g., *Pseudomonas* spp. and *Streptomyces* spp.) across the crop diversity gradient; however, we did not detect distinct changes in putative biocontrol community composition (Appendix S1: Table S4).

Our study revealed that cover crops in combination with corn-soy-wheat rotations increased abundance of the *prnD* gene, which is responsible for producing antifungal compound pyrrolnitrin (PRN) (Garbeva et al. 2004b, Haas and Défago 2005), by about 9 % compared to the other cropping system. Cover crop species may have important effects on the *prnD* gene abundance and disease suppressive functional potential in soils, but only in combination with corn-soy-wheat because the cover crop with corn only did not show high *prnD* abundance (Fig. 3). The *prnD* gene abundance in all cropping systems was higher than in fallow treatment (or most diverse treatment). The abundance of DAPG and PRN producers increasing with plant diversity has been previously observed (Latz et al. 2012). Compared to agricultural soils, the PRN producers were more frequently detected in grassland or grassland-derived plots (Garbeva et al. 2004a, 2004b). In a previous study, the *prnD* gene abundance increased in the presence of grasses, but the legume species tended to decrease the DAPG and PRN producer abundance (Latz et al. 2012). Without crops (as reflected in the fallow treatment), we observed that disease suppressive potential significantly declined. This observation may be indicative of species-specific facilitation of PPS soil microorganisms. This disease suppressive phenomenon is known to have important implications for sustainable biocontrol of soil-borne pathogens. In addition, it is possible that when plant diversity is high, there is less soil-borne pathogen pressure on plant hosts due to decreased competition for resources among pathogenic and non-pathogenic soil microorganisms.

#### Proposed mechanisms for crop diversity effects on soil bacterial diversity and PPS abundance

Disease suppression may have a major role in what is colloquially referred to as “the rotation effect.” Our study provided evidence that crop diversity alters soil bacterial community composition and population of PPS microbes, but the mechanisms through which this occurs can include physical, chemical, and biological changes to the soil environment. Crops can influence soil properties and soil microbiomes in a variety of ways, including physically and chemically. Cover crops are the most salient feature of these crop rotations affecting the soil bacterial community in general. This is not surprising, since cover crops have been shown to influence several soil properties, which likely have indirect effects on the soil bacterial community composition. In addition, previous studies showed cover crops can have immediate impacts on soil microbial communities (Wiggins and Kinkel 2005, Finney et al. 2017). Soil properties like total C, total N, pH, and bulk density and porosity have all been shown to increase with cover crops (Bullock 1992, Liebman and Dyck 1993, Tilman et al. 2002, McDaniel et al. 2014b, Tiemann et al. 2015). Physically, crop diversity (especially rotations) can enhance soil properties like improving plant water availability by lowering bulk density, increasing soil pore space, and increasing soil aggregate formation (Tilman et al. 2002, McDaniel et al. 2014b, Tiemann et al. 2015), which could have indirect influence over the soil bacterial community as well. Chemically, cover crops are providing more carbon to the soil through residues, but also root exudation of recently assimilated photosynthate, composed of soluble, low molecular weight organic compounds (Neumann and Romheld 2007). As a consequence, the increased C flow from cover crop root exudates can stimulate soil microbial activity. Changes in root exudates have been observed to shift microbial community composition and stimulate a diverse microbial community (Hooper et al. 2000, Stephan et al. 2000, Paterson et al. 2009, Dijkstra et al. 2010). Biologically, some soil microorganisms can provide PPS through competition for nutrients, antibiosis, and induction of system resistance of host plants (Doornbos et al. 2012). Our study focused on soil bacterial community composition. It has been identified that crop rotation also influences soil fungal and faunal communities, which are also important members of the soil food web (McLaughlin and Mineau 1995). For example, increased protist predation on soil bacteria has resulted in indirect effect on disease suppression function (Jousset et al. 2008, 2010). These studies revealed that increased predator pressure by soil protists have been linked to increased biocontrol function through enhanced bacterial DAPG production (Jousset et al. 2008, 2010).

However, disease suppression traits such as antifungal production may not be needed and are not maintained in the community when crops are no longer planted. Several explanations could underpin our observations. When agricultural management is absent, there is reduced selection for soil microorganisms with disease suppression traits. Higher plant diversity reflected in longer crop rotations was expected to support overall diversity, resulting in an increased probability of getting more disease suppressive microbes. However, we observed that overall taxonomic diversity decreased with increasing crop diversity, indicating alternative mechanisms maybe be involved in diversity-function relationship. One argument is that in monocultures, the selection for fungal pathogen defense is weakened and microbes that are (constitutively or facultatively) making defense compounds are paying a cost and are replaced by microbes that do not invest in the defense strategy. In addition, fluctuating environments can influence selection of traits (Heath et al. 2010, Akçay and Simms 2011). For example, high variation in carbon compounds such as under diverse crop rotations could alter selection of defense traits, whereby crop plants facilitate PPS or other defense traits that are adaptive only when crop plants are present. Increasing plant diversity such as in fallow, non-cropping systems, provides opportunity for microbial community members to partition according to diverse (and more even) carbon resources rather than crop inputs driving selection of microbial communities and defense traits (Hartmann et al. 2009). In other words, when you are in a resource rich soil under fallow, there no need for PPS gene production and maintenance. Our findings combined with previous studies suggest that the land-use regime, plant diversity, and plant species influence disease suppressive microbial communities.

## Conclusions

We and others demonstrate links between crop diversity soil ecosystem functions; however, the mechanisms underpinning this relationship require further study for more predictive soil microbiome management (Lauber et al. 2008, Jangid et al. 2008, McDaniel et al. 2014b, Orr et al. 2015, Tiemann et al. 2015, Venter et al. 2016). Crop diversity may facilitate the abundance of PPS organisms even though both our study and previous study show decreases in structural diversity and functional evenness (McDaniel and Grandy 2016). We observed that the soil microbial community composition may be more important than soil microbial diversity to soil disease suppression. Crop rotations may also provide other important benefits like enhanced nutrient provisioning to plants, improvement of soil physical properties, increases in soil C, and increases in soil microbial and faunal activity that also could be responsible for the increased yields responsible for the rotation effect (Ball et al. 2005, van der Putten et al. 2016). Additional research focused on identifying patterns in soil microbial diversity and ecosystem function relationships can inform microbiome management, which will involve defined management of soil nutrients and plant diversity.

## Acknowledgements

This work was supported by the U.S. Department of Agriculture National Institute of Food and Agriculture Postdoctoral Fellowship (2012-67012-19845 to A.L.P.) and the National Science Foundation (DEB 1442246 to J.T.L.). The funders had no role in study design, data collection and interpretation, preparation of the manuscript, or decision to submit the work for publication. Support was also provided by the NSF Long-term Ecological Research Program (DEB 1027253) at the Kellogg Biological Station and by Michigan State University AgBioResearch. We would like to thank the Kellogg Biological Station LTER for logistical support and use of sampling sites. We also acknowledge the logistical support of K.L. Gross and G.P. Robertson, who originally established these sites. We also thank M. Muscarella, J. Ford, S. Krahnke, and M. Brewer for microbial analyses support and B. O’Neill, A.S. Grandy and T.M. Schmidt Labs for field and soil analyses support. All code and data used in this study can be found in a public GitHub repository (https://github.com/PeraltaLab/CropDiversity) and the NCBI SRA (BioProject).

## Literature Cited

Akçay, E., and E. L. Simms. 2011. Negotiation, sanctions, and context dependency in the legume-rhizobium mutualism. The American Naturalist 178:1–14.

Allison, S. D., S. S. Chacon, and D. P. German. 2014. Substrate concentration constraints on microbial decomposition. Soil Biology and Biochemistry 79:43–49.

Badri, D. V., N. Quintana, E. G. E. Kassis, H. K. Kim, Y. H. Choi, A. Sugiyama, R. Verpoorte, E. Martinoia, D. K. Manter, and J. M. Vivanco. 2009. An ABC transporter mutation alters root exudation of phytochemicals that provoke an overhaul of natural soil microbiota. Plant Physiology 151:2006–2017.

Bakker, M. G., J. D. Glover, J. G. Mai, and L. L. Kinkel. 2010. Plant community effects on the diversity and pathogen suppressive activity of soil streptomycetes. Applied Soil Ecology 46:35–42.

Ball, B. C., I. Bingham, R. M. Rees, C. A. Watson, and A. Litterick. 2005. The role of crop rotations in determining soil structure and crop growth conditions. Canadian Journal of Soil Science 85:557–577.

Berendsen, R. L., C. M. J. Pieterse, and P. A. H. M. Bakker. 2012. The rhizosphere microbiome and plant health. Trends in Plant Science 17:478–486.

Berg, G., and K. Smalla. 2009. Plant species and soil type cooperatively shape the structure and function of microbial communities in the rhizosphere. FEMS Microbiology Ecology 68:1–13.

Bever, J. D., K. M. Westover, and J. Antonovics. 1997. Incorporating the soil community into plant population dynamics: the utility of the feedback approach. Journal of Ecology 85:561–573.

Brussaard, L., P. C. de Ruiter, and G. G. Brown. 2007. Soil biodiversity for agricultural sustainability. Agriculture, Ecosystems & Environment 121:233–244.

Bullock, D. G. 1992. Crop rotation. Critical Reviews in Plant Sciences 11:309–326.

Caporaso, J. G., C. L. Lauber, W. A. Walters, D. Berg-Lyons, J. Huntley, N. Fierer, S. M. Owens, J. Betley, L. Fraser, M. Bauer, N. Gormley, J. A. Gilbert, G. Smith, and R. Knight. 2012. Ultra-high-throughput microbial community analysis on the Illumina HiSeq and MiSeq platforms. The ISME Journal 6:ismej20128.

Chaparro, J. M., A. M. Sheflin, D. K. Manter, and J. M. Vivanco. 2012. Manipulating the soil microbiome to increase soil health and plant fertility. Biology and Fertility of Soils 48:489–499.

Clarke, K. R. 1993. Non-parametric multivariate analyses of changes in community structure. Australian Journal of Ecology 18:117–143.

Crum, J. R., and H. P. Collins. 1995. KBS Soils [Online]. www.lter.kbs.msu.edu/soil/characterization.

Culman, S. W., S. S. Snapp, M. A. Freeman, M. E. Schipanski, J. Beniston, R. Lal, L. E. Drinkwater, A. J. Franzluebbers, J. D. Glover, A. S. Grandy, J. Lee, J. Six, J. E. Maul, S. B. Mirksy, J. T. Spargo, and M. M. Wander. 2012. Permanganate oxidizable carbon reflects a processed soil fraction that is sensitive to management. Soil Science Society of America Journal 76:494–504.

Dijkstra, F. A., J. A. Morgan, D. Blumenthal, and R. F. Follett. 2010. Water limitation and plant inter-specific competition reduce rhizosphere-induced C decomposition and plant N uptake. Soil Biology and Biochemistry 42:1073–1082.

Doornbos, R. F., L. C. van Loon, and P. A. H. M. Bakker. 2012. Impact of root exudates and plant defense signaling on bacterial communities in the rhizosphere. A review. Agronomy for Sustainable Development 32:227–243.

Edgar, R. C., B. J. Haas, J. C. Clemente, C. Quince, and R. Knight. 2011. UCHIME improves sensitivity and speed of chimera detection. Bioinformatics 27:2194–2200.

von Felten, A., J. B. Meyer, G. Défago, and M. Maurhofer. 2011. Novel T-RFLP method to investigate six main groups of 2, 4-diacetylphloroglucinol-producing pseudomonads in environmental samples. Journal of Microbiological Methods 84:379–387.

Finney, D. M., J. S. Buyer, and J. P. Kaye. 2017. Living cover crops have immediate impacts on soil microbial community structure and function. Journal of Soil and Water Conservation 72:361–373.

Garbeva, P., J. Postma, J. A. Van Veen, and J. D. Van Elsas. 2006. Effect of above-ground plant species on soil microbial community structure and its impact on suppression of Rhizoctonia solani AG3. Environmental Microbiology 8:233–246.

Garbeva, P., J. A. van Veen, and J. D. van Elsas. 2004a. Microbial diversity in soil: Selection of microbial populations by plant and soil type and implications for disease suppressiveness. Annual Review of Phytopathology 42:243–270.

Garbeva, P., K. Voesenek, and J. D. van Elsas. 2004b. Quantitative detection and diversity of the pyrrolnitrin biosynthetic locus in soil under different treatments. Soil Biology and Biochemistry 36:1453–1463.

Goldfarb, K. C., U. Karaoz, C. A. Hanson, C. A. Santee, M. A. Bradford, K. K. Treseder, M. D. Wallenstein, and E. L. Brodie. 2011. Differential growth responses of soil bacterial taxa to carbon substrates of varying chemical recalcitrance. Frontiers in Microbiology 2.

Haas, D., and G. Défago. 2005. Biological control of soil-borne pathogens by fluorescent pseudomonads. Nature Reviews Microbiology 3:nrmicro1129.

Hartmann, A., M. Schmid, D. van Tuinen, and G. Berg. 2009. Plant-driven selection of microbes. Plant and Soil 321:235–257.

Heath, K. D., A. J. Stock, and J. R. Stinchcombe. 2010. Mutualism variation in the nodulation response to nitrate. Journal of Evolutionary Biology 23:2494–2500.

Hooper, D. U., D. E. Bignell, V. K. Brown, L. Brussard, J. M. Dangerfield, D. H. Wall, D. A. Wardle, D. C. Coleman, K. E. Giller, P. Lavelle, V. D. Putten, W. H. D. Ruiter, P. C. J. Rusek, W. L. Silver, J. M. Tiedje, and V. Wolters. 2000. Interactions between aboveground and belowground biodiversity in terrestrial ecosystems: patterns, mechanisms, and feedbacks. BioScience 50:1049–1061.

Jangid, K., M. A. Williams, A. J. Franzluebbers, J. S. Sanderlin, J. H. Reeves, M. B. Jenkins, D. M. Endale, D. C. Coleman, and W. B. Whitman. 2008. Relative impacts of land-use, management intensity and fertilization upon soil microbial community structure in agricultural systems. Soil Biology and Biochemistry 40:2843–2853.

Jousset, A., L. Rochat, S. Scheu, M. Bonkowski, and C. Keel. 2010. Predator-prey chemical warfare determines the expression of biocontrol genes by rhizosphere-associated pseudomonas fluorescens. Applied and Environmental Microbiology 76:5263–5268.

Jousset, A., S. Scheu, and M. Bonkowski. 2008. Secondary metabolite production facilitates establishment of rhizobacteria by reducing both protozoan predation and the competitive effects of indigenous bacteria. Functional Ecology 22:714–719.

Karlen, D. L., G. E. Varvel, D. G. Bullock, and R. M. Cruse. 1994. Crop Rotations for the 21st Century. Pages 1–45 in D. L. Sparks, editor. Advances in Agronomy. Academic Press.

Kozich, J. J., S. L. Westcott, N. T. Baxter, S. K. Highlander, and P. D. Schloss. 2013. Development of a dual-index sequencing strategy and curation pipeline for analyzing amplicon sequence data on the MiSeq Illumina sequencing platform. Applied and Environmental Microbiology:AEM.01043–13.

Kulmatiski, A., and K. H. Beard. 2011. Long-term plant growth legacies overwhelm short-term plant growth effects on soil microbial community structure. Soil Biology and Biochemistry 43:823–830.

Kulmatiski, A., K. H. Beard, J. R. Stevens, and S. M. Cobbold. 2008. Plant–soil feedbacks: a meta-analytical review. Ecology Letters 11:980–992.

Latz, E., N. Eisenhauer, B. C. Rall, E. Allan, C. Roscher, S. Scheu, and A. Jousset. 2012. Plant diversity improves protection against soil-borne pathogens by fostering antagonistic bacterial communities. Journal of Ecology 100:597–604.

Lau, J. A., and J. T. Lennon. 2012. Rapid responses of soil microorganisms improve plant fitness in novel environments. Proceedings of the National Academy of Sciences 109:14058–14062.

Lauber, C. L., M. S. Strickland, M. A. Bradford, and N. Fierer. 2008. The influence of soil properties on the structure of bacterial and fungal communities across land-use types. Soil Biology and Biochemistry 40:2407–2415.

Liebman, M., and E. Dyck. 1993. Crop rotation and intercropping strategies for weed management. Ecological Applications 3:92–122.

Lin, B. B. 2011. Resilience in agriculture through crop diversification: adaptive management for environmental change. BioScience 61:183–193.

Lugtenberg, B., and F. Kamilova. 2009. Plant-growth-promoting rhizobacteria. Annual Review of Microbiology 63:541–556.

Mavrodi, O. V., D. V. Mavrodi, J. A. Parejko, L. S. Thomashow, and D. M. Weller. 2012. Irrigation differentially impacts populations of indigenous antibiotic-producing pseudomonas spp. in the rhizosphere of wheat. Applied and Environmental Microbiology 78:3214–3220.

McDaniel, M. D., and A. S. Grandy. 2016. Soil microbial biomass and function are altered by 12 years of crop rotation. SOIL 2:583–599.

McDaniel, M. D., A. S. Grandy, L. K. Tiemann, and M. N. Weintraub. 2014a. Crop rotation complexity regulates the decomposition of high and low quality residues. Soil Biology and Biochemistry 78:243–254.

McDaniel, M. D., L. K. Tiemann, and A. S. Grandy. 2014b. Does agricultural crop diversity enhance soil microbial biomass and organic matter dynamics? A meta-analysis. Ecological Applications 24:560–570.

McLaughlin, A., and P. Mineau. 1995. The impact of agricultural practices on biodiversity. Agriculture, Ecosystems & Environment 55:201–212.

Mendes, L. W., S. M. Tsai, A. A. Navarrete, M. de Hollander, J. A. van Veen, and E. E. Kuramae. 2015. Soil-borne microbiome: linking diversity to function. Microbial Ecology 70:255–265.

Mendes, R., P. Garbeva, and J. M. Raaijmakers. 2013. The rhizosphere microbiome: significance of plant beneficial, plant pathogenic, and human pathogenic microorganisms. FEMS Microbiology Reviews 37:634–663.

Mills, K. E., and J. D. Bever. 1998. Maintenance of diversity within plant communities: soil pathogens as agents of negative feedback. Ecology 79:1595–1601.

Muscarella, M., K. Bird, M. Larsen, S. Placella, and J. Lennon. 2014. Phosphorus resource heterogeneity in microbial food webs. Aquatic Microbial Ecology 73:259–272.

Naeem, S., and J. P. Wright. 2003. Disentangling biodiversity effects on ecosystem functioning: deriving solutions to a seemingly insurmountable problem. Ecology Letters 6:567–579.

Needleman, S. B., and C. D. Wunsch. 1970. A general method applicable to the search for similarities in the amino acid sequence of two proteins. Journal of Molecular Biology 48:443–453.

Neumann, G., and V. Romheld. 2007. The release of root exudates as affects by the plant physiological status. Pages 23–72 in R. Pinton, Z. Varanini, and P. Nannipieri, editors. The Rhizosphere: Biochemistry and Organic Substances at the Soil-plant Interface. 2nd edition. CRC Press.

Oksanen, J., F. G. Blanchet, R. Kindt, P. Legendre, R. B. O’Hara, G. L. Simpson, P. Solymos, M. H. H. Stevens, and H. Wagner. 2010. Community Ecology Package “vegan.”

Orr, C. h., C. j. Stewart, C. Leifert, J. m. Cooper, and S. p. Cummings. 2015. Effect of crop management and sample year on abundance of soil bacterial communities in organic and conventional cropping systems. Journal of Applied Microbiology 119:208–214.

Packer, A., and K. Clay. 2000. Soil pathogens and spatial patterns of seedling mortality in a temperate tree. Nature 404:278–281.

Paterson, E., A. J. Midwood, and P. Millard. 2009. Through the eye of the needle: a review of isotope approaches to quantify microbial processes mediating soil carbon balance. New Phytologist 184:19–33.

Penton, C. R., V. V. S. R. Gupta, J. M. Tiedje, S. M. Neate, K. Ophel-Keller, M. Gillings, P. Harvey, A. Pham, and D. K. Roget. 2014. Fungal community structure in disease suppressive soils assessed by 28s lsu gene sequencing. PLOS ONE 9:e93893.

Peralta, A. L., J. W. Matthews, D. N. Flanagan, and A. D. Kent. 2012. Environmental factors at dissimilar spatial scales influence plant and microbial communities in restored wetlands. Wetlands 32:1125–1134.

Postma, J., M. T. Schilder, J. Bloem, and W. K. van Leeuwen-Haagsma. 2008. Soil suppressiveness and functional diversity of the soil microflora in organic farming systems. Soil Biology and Biochemistry 40:2394–2406.

van der Putten, W. H., R. D. Bardgett, J. D. Bever, T. M. Bezemer, B. B. Casper, T. Fukami, P. Kardol, J. N. Klironomos, A. Kulmatiski, J. A. Schweitzer, K. N. Suding, T. F. J. Van de Voorde, and D. A. Wardle. 2013. Plant–soil feedbacks: the past, the present and future challenges. Journal of Ecology 101:265–276.

van der Putten, W. H., M. A. Bradford, E. Pernilla Brinkman, T. F. J. van de Voorde, and G. F. Veen. 2016. Where, when and how plant–soil feedback matters in a changing world. Functional Ecology 30:1109–1121.

Raaijmakers, J. M., T. C. Paulitz, C. Steinberg, C. Alabouvette, and Y. Moënne-Loccoz. 2009. The rhizosphere: a playground and battlefield for soilborne pathogens and beneficial microorganisms. Plant and Soil 321:341–361.

Reardon, C. L., H. T. Gollany, and S. B. Wuest. 2014. Diazotroph community structure and abundance in wheat–fallow and wheat–pea crop rotations. Soil Biology and Biochemistry 69:406–412.

Robertson, G. P., and S. K. Hamilton. 2015. Long-term ecological research in agricultural landscapes at the Kellogg Biological Station LTER site: conceptual and experimental framework. Page 1–32 in S. K. Hamilton, J. E. Doll, and G. P. Robertson, editors. The Ecology of Agricultural Landscapes: Long-Term Research on the Path to Sustainability. Oxford University Press, New York, New York, USA.

Schlatter, D., L. Kinkel, L. Thomashow, D. Weller, and T. Paulitz. 2017. Disease suppressive soils: new insights from the soil microbiome. Phytopathology 107:1284–1297.

Schloss, P. D., S. L. Westcott, T. Ryabin, J. R. Hall, M. Hartmann, E. B. Hollister, R. A. Lesniewski, B. B. Oakley, D. H. Parks, C. J. Robinson, J. W. Sahl, B. Stres, G. G. Thallinger, D. J. V. Horn, and C. F. Weber. 2009. Introducing mothur: open-source, platform-independent, community-supported software for describing and comparing microbial communities. Applied and Environmental Microbiology 75:7537–7541.

Smith, R. G., and K. L. Gross. 2006. Weed community and corn yield variability in diverse management systems. Weed Science 54:106–113.

Smith, R. G., and K. L. Gross. 2007. Assembly of weed communities along a crop diversity gradient. Journal of Applied Ecology 44:1046–1056.

Smukler, S. M., S. Sánchez-Moreno, S. J. Fonte, H. Ferris, K. Klonsky, A. T. O’Geen, K. M. Scow, K. L. Steenwerth, and L. E. Jackson. 2010. Biodiversity and multiple ecosystem functions in an organic farmscape. Agriculture, Ecosystems & Environment 139:80–97.

Stephan, A., A. H. Meyer, and B. Schmid. 2000. Plant diversity affects culturable soil bacteria in experimental grassland communities. Journal of Ecology 88:988–998.

Tiemann, L. K., A. S. Grandy, E. E. Atkinson, E. Marin-Spiotta, and M. D. McDaniel. 2015. Crop rotational diversity enhances belowground communities and functions in an agroecosystem. Ecology Letters 18:761–771.

Tilman, D., C. Balzer, J. Hill, and B. L. Befort. 2011. Global food demand and the sustainable intensification of agriculture. Proceedings of the National Academy of Sciences 108:20260–20264.

Tilman, D., K. G. Cassman, P. A. Matson, R. Naylor, and S. Polasky. 2002. Agricultural sustainability and intensive production practices. Nature 418:nature01014.

Venter, Z. S., K. Jacobs, and H.-J. Hawkins. 2016. The impact of crop rotation on soil microbial diversity: A meta-analysis. Pedobiologia 59:215–223.

Warton, D. I., S. T. Wright, and Y. Wang. 2012. Distance-based multivariate analyses confound location and dispersion effects. Methods in Ecology and Evolution 3:89–101.

Weller, D. M., J. M. Raaijmakers, B. B. M. Gardener, and L. S. Thomashow. 2002. Microbial Populations Responsible for Specific Soil Suppressiveness to Plant Pathogens. Annual Review of Phytopathology 40:309–348.

Wiggins, B. E., and L. L. Kinkel. 2005. Green manures and crop sequences influence alfalfa root rot and pathogen inhibitory activity among soil-borne streptomycetes. Plant and Soil 268:271–283.

Yilmaz, P., L. W. Parfrey, P. Yarza, J. Gerken, E. Pruesse, C. Quast, T. Schweer, J. Peplies, W. Ludwig, and F. O. Glöckner. 2014. The SILVA and “All-species Living Tree Project (LTP)” taxonomic frameworks. Nucleic Acids Research 42:D643–D648.

Yin, C., K. L. Jones, D. E. Peterson, K. A. Garrett, S. H. Hulbert, and T. C. Paulitz. 2010. Members of soil bacterial communities sensitive to tillage and crop rotation. Soil Biology and Biochemistry 42:2111–2118.

Zak, D. R., W. E. Holmes, D. C. White, A. D. Peacock, and D. Tilman. 2003. Plant diversity, soil microbial communities, and ecosystem function: are there any links? Ecology 84:2042–2050.

